# Evaluation of Parallel Accumulation-Serial Fragmentation methods for metaproteomics using a model microbiome

**DOI:** 10.1101/2025.08.13.670166

**Authors:** Ruben Shreshta, Andrew T. Rajczewski, Aleksey Ogurtsov, Katherine T. Do, Gelio Alves, Yi-Kuo Yu, Matthew Willets, Manuel Kleiner, Timothy J. Griffin, Pratik D. Jagtap

## Abstract

We present a multi-organism, ground-truth metaproteomic dataset generated using data-dependent acquisition Parallel Accumulation-Serial Fragmentation (ddaPASEF) and data-independent acquisition (diaPASEF) on a trapped ion mobility mass spectrometer. The dataset comprises 28 species (30 strains) spanning Archaea, Bacteria, Eukarya, and bacteriophages, assembled into an uneven mock community with a 400-fold dynamic range of organismal abundance. Four biological replicates were processed using a standardized sample preparation workflow and analyzed using described LC-MS conditions. Raw data, search outputs, protein coverage metrics, quantitative abundance estimates, and strain-specific peptide evidence are provided. The dataset enables evaluation of proteome depth, proteome coverage, quantitative reproducibility, and strain-level resolution across acquisition strategies. diaPASEF yielded broader peptide and protein detection and deeper sequence coverage, while both methods accurately quantified high-abundance organisms. Detection evidence for low-abundance bacteriophages and single-amino-acid-variant peptides is included to support benchmarking of taxonomic and strain-resolved analyses. This dataset provides a comprehensive reference for developing and validating metaproteomic workflows, protein inference strategies, and ion mobility-aware computational tools.

## BACKGROUND AND SUMMARY

Mass spectrometry(MS)-based metaproteomics allows for the identification and quantification of thousands of proteins from clinical and environmental samples and is rapidly gaining importance in microbiome sciences^1,2^. Metaproteomics enables direct measurement of proteins expressed by complex microbial communities, providing taxonomic and functional information that complements metagenomics and metabolomics^3^. Measuring functional response through characterization of microbiome-expressed proteins, has the potential to shed light on the mechanistic details of how a microbiome interacts with its immediate environment^4^.

For sensitive and accurate MS-based metaproteomics, several challenges must be overcome. The metaproteome encompasses a vast dynamic abundance range, including numerous low-abundance proteins that are often undetectable with conventional mass spectrometry methods. Moreover, the complexity at taxonomic level and protein inference makes robust identification and accurate quantification challenging. MS-based metaproteomics commonly employs Data-Dependent Acquisition (DDA) and Data-Independent Acquisition (DIA), two distinct strategies that differ in peptide selection and coverage^5-6^. While DDA targets the most abundant ions in real time, often missing low-abundance peptides, DIA fragments all ions within set m/z windows, offering broader and more reproducible coverage at the cost of increased data complexity^5-6^. The diaPASEF approach has the potential to overcome these limitations by integrating trapped ion mobility spectrometry (TIMS) with Parallel Accumulation–Serial Fragmentation (PASEF), which introduces an additional ion mobility separation dimension. This enables the resolution of co-eluting and isobaric peptides, significantly improving the confidence and accuracy of peptide identification and quantification. PASEF achieves a 100% duty cycle by accumulating ions in parallel and releasing them sequentially for fragmentation, thereby maximizing ion utilization and increasing sequencing speed without compromising sensitivity. As a result, diaPASEF provides deep and comprehensive coverage of both microbial (and host) proteomes within a single, Liquid Chromatography (LC)-MS run. Recently, metaproteomics researchers have used diaPASEF for clinical metaproteomics studies^7-11^.

As metaproteomics continues to evolve, a growing community of researchers is developing increasingly sophisticated software tools^12-20^ to analyze complex microbial protein data. However, progress is often hindered by the lack of high-quality, ground-truth datasets that can serve as reliable benchmarks for tool evaluation and comparison. In this study, we used a mock community containing a diversity of microbes from all three domains of life^21^, as well as bacteriophages to assess the benefits of using a ddaPASEF and diaPASEF method for quantitative metaproteomics analysis. The dataset includes raw mass spectrometry files, database search results, peptide and protein identifications, protein coverage metrics, quantitative abundance estimates, and strain-specific peptide evidence. These data support benchmarking of acquisition depth, reproducibility, protein inference, strain discrimination, and quantitative performance. The inclusion of bacteriophage peptides and low-abundance taxa provides a realistic test for workflows targeting rare organisms. This dataset serves as a resource for developing next-generation metaproteomics tools, including DIA-specific algorithms, mobility-aware scoring methods, and improved taxonomic assignment strategies.

## METHODS

### Samples

We extracted four biological replicates of a ground-truth mock community containing Archaea, Bacteria, Eukarya and Bacteriophages with known species abundances^21^. The uneven mock community used in this study was designed to encompass a broad range of bacterial phylogeny (**Supplementary Table 1**) and accommodate varying species abundances, aiming to evaluate the dynamic range and detection limits (**Figure 1**).

**Figure 1:**
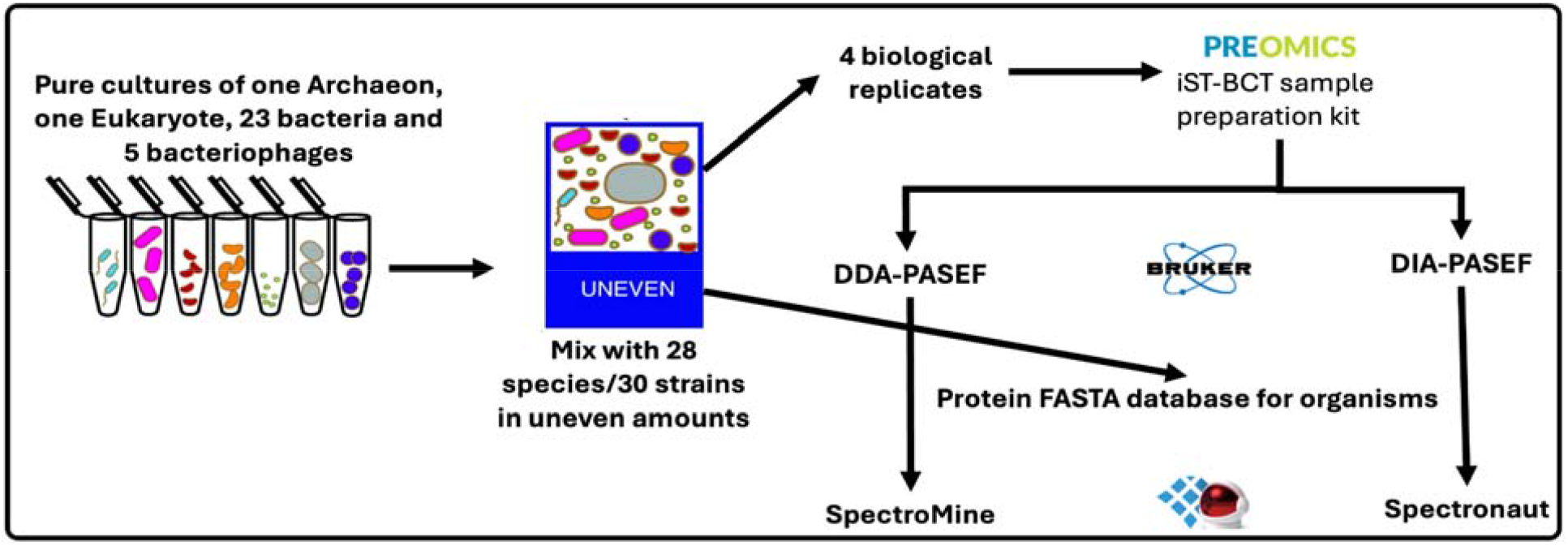
Overview of experimental design and dataset generation. This study used a ground-truth sample comprising 28 species (30 strains) of Archaea, Bacteria, Eukarya, and bacteriophages with known species abundances. These species were used to assemble an uneven mock community, wherein species abundances varied at both cell number and proteinaceous biomass levels^21^. Four biological replicates of this uneven mock community were processed using the PreOmics iST-BCT kit. These samples were analyzed using LC-MS and ddaPASEF or diaPASEF mass spectrometry. This ddaPASEF and diaPASEF mass spectrometry data was searched against a protein sequence database (containing the proteomes belonging to all species in the uneven mock community) using SpectroMine and Spectronaut, respectively.

### Sample preparation

Four biological replicates for the uneven mock community samples were processed using the PreOmics iST-BCT kit (**Figure 1**), which is used for processing of biological fluid samples for proteomics analysis using mass spectrometry^22^. The kit includes all necessary reagents for lysis, reduction, and alkylation using the LYSE reagent at 95°C for 10 minutes, followed by digestion with the DIGEST reagent at 37°C for 60 minutes. The peptide-digest is cleaned up through cartridge processing, including washing, elution, drying, and resuspending with the LOAD reagent, and ready for mass spectrometry analysis.

### Sample processing

Dried peptide samples were resuspended in 10 uL of 3% acetonitrile (ACN), 0.1% formic acid (FA) and vortexed and sonicated for 30s; centrifuged and transferred to vials. 750 ng of peptide digest was loaded for ddaPASEF and 1 ug of sample was loaded for diaPASEF. We used four replicates of mock community types with uneven amounts of protein from each organism (**Figure 1** and **Supplementary Table 1**).

### LC-MS methods

Peptides were reconstituted in 3% ACN/0.1% FA then analyzed by liquid chromatography-mass spectrometry (LC-MS), with the following conditions: Nanoflow reversed-phase liquid chromatography was performed on a nanoElute system (Bruker Daltonics) connected to a hybrid TIMS QTOF mass spectrometer (Bruker timsTOF HT) via a CaptiveSpray nano-electrospray source. Peptides were separated using an 80- and 120-min gradient on a C18 column (25cm × 75 μm, 1.7 μm, IonOpticks). The mass spectrometer was operated in ddaPASEF and diaPASEF mode. For ddaPASEF analysis, the mobile phase A was water with 0.1 vol% FA and mobile phase B was ACN with 0.1 vol% FA. Peptides were separated with a linear and stepped gradient ranging from 2 to 28 solvent B over 85 min, 28 to 32% in 25 min, 32 to 85% in 0.1 min, and held at 85% for 9.9 min at 300 nl/min. MS scans were acquired from 100 to 1700 *m/z* and 1/K0□=□0.6 Vs cm−2 to 1.6 Vs cm−2. Ten PASEF ramps were acquired with an accumulation and ramp time of 100 ms. Precursors above the minimum intensity threshold of 1000 were isolated with 2 Th at□<□700 *m/z* or 3 Th >800 *m/z* and reacquired until a target intensity of 10,000. Active exclusion was set to 40 s, and the collision energy (CE) was a linear function between the CE and the ion-mobility starting from 20 eV at 1/K0□=□0.6 Vs cm-2 to 59 eV at 1/K0□=□1.6 V cm-2. For diaPASEF analysis, peptides were separated with a linear and stepped gradient ranging from 2 to 28 solvent B over 63 min, 28 to 32% in 7 min, 32 to 85% in 0.1 min, and held at 85% for 9.9 min at 300 nl/min. MS scans were acquired from 100 to 1700 *m/z* and 1/K0 = 0.7 Vs cm-2 to 1.3 Vs cm-2. The diaPASEF method can be found in **Supplementary Table 2**, that covered the range of 300 to 1200m/z and an ion mobility range of 0.7-1.33 Vs cm-2, with 35Da isolation windows (1Da overlap) with a total cycle time of 850 ms. The collision energy (CE) was a linear function between the CE and the ion-mobility starting from 20 eV at 1/K0□= 0.6 Vs cm-2 to 59 eV at 1/K0 = 1.6 V cm-2. Accumulation and ramp time were set to 100 ms.

### Data analysis

A protein sequence database for the uneven mock community was generated by combining the proteomes of all species in the mock community into a FASTA file called “Mock_Comm_RefDB_V3_Clustered95.fasta” (112,580 sequences). Proteomes were acquired from UniProtKB or NCBI and are detailed in **Supplementary Table 1**.

### ddaPASEF search in SpectroMine

ddaPASEF mass spectrometry data were searched against the customized mock community protein sequence database with SpectroMine v.4 (Biognosys) with default Pulsar search settings. In brief, each run was assigned to its own block; Trypsin/P as specific enzyme; peptide length from 7 to 52; max missed cleavages 2; toggle N-terminal M turned on; maximum of five variable modifications; Carbamidomethyl on C as fixed modification; Oxidation on M and Acetyl at protein N-terminus as variable modifications; FDRs at PSM, peptide and protein level all set to 0.01.

### DirectDIA search in Spectronaut

diaPASEF mass spectrometry data were processed in Spectronaut v.18 (Biognosys) using the library-free directDIA mode supplemented with hybrid library extension from ddaPASEF runs. The default search parameters were used along with the customized protein sequence database for the mock community. In brief, the settings for Pulsar were the same as mentioned above for ddaPASEF Spectromine search. diaPASEF Analysis settings were: 0.01 Q value cutoff at precursor, peptide and protein level. Peptide precursors are calculated at the peptide ion level, where each unique combination of peptide sequence with modifications and charge state is considered a distinct precursor. For example, a peptide observed in both the 2+ and 3+ charge states contributes two peptide precursor identifications. Peptide counts, however, are calculated at the sequence level with modifications, such that all charge states corresponding to the same peptide sequence are consolidated into a single peptide identification. LFQ quantitation at MS2 level using area as quantitation type with cross-run normalization enabled. Spectronaut uses the IDPicker algorithm^23^ for protein inference.

### Protein Coverage Analysis

Protein coverage was calculated to assess how comprehensively species-specific proteins were detected in ddaPASEF and diaPASEF approaches. For each organism in the ground-truth sample, all proteins uniquely assigned to that species (species-specific proteins) were extracted, and the percent sequence coverage for each protein was obtained from the corresponding search results. Coverage values represent the proportion of amino acids within a protein sequence that were matched by confidently identified peptides. We selected proteins that were detected by both ddAPASEF and diaPASEF datasets and for each organism, the average protein coverage was computed separately for ddaPASEF and diaPASEF datasets by taking the mean coverage across all species-specific proteins.

### Single amino acid variant peptides and strain detection analysis

Theoretical peptides were identified from the reference proteomes of each strain pair (*Staphylococcus aureus* ATCC 13709 vs. ATCC 25923; *Rhizobium leguminosarum* bv. *viciae* 3841 vs. VF39). These theoretical peptides were matched with the detected peptides in both the ddaPASEF and diaPASEF datasets. For this, the searches were performed against a database with all four strains added without 95% clustering. Peptides unique to one strain or differing by a single amino acid substitution between strains were identified, and single amino acid variant (SAV) peptide pairs selected for analysis are listed in **Supplementary Table 3** for diaPASEF data and **Supplementary Table 4** for ddaPASEF data.

Peptides were then grouped according to their strain specificity: detected exclusively in one strain (“Only 137” or “Only 841”), exclusively in the other strain (“Only 259” or “Only VF”), or detected in both strains (“Both”). Analyses were performed independently for ddaPASEF and diaPASEF datasets.

### Quantitative Analysis

For peptide quantification, peptide intensities from proteins of each organism were summed. The abundance of each organism was represented as the sum intensity of all unique peptides annotated to that organism. The percentage of the total peptide intensity was then compared to the expected percentage for each organism. Species abundances were calculated for each acquisition method (ddaPASEF or diaPASEF) by summing the unique peptide intensities of individual species and dividing by the total peptide intensity across all species within each replicate. These values were then averaged to determine the mean percentage abundances. The summed peptide intensities for each abundance tier were compared to their expected percentages. Peptides unambiguously assigned to a single organism were used to calculate species abundances within the microbiome community, following the methodology outlined in the original study for which these samples were generated^21^.

## DATA RECORDS

All data are available through the MassIVE repository under accession **MSV000103357**. The dataset includes: a) Raw LC-MS files (ddaPASEF and diaPASEF) in zipped .d folders, b) Search results (SpectroMine and Spectronaut) c) Peptide identification tables, c) Protein sequence FASTA databases, d) Supplementary figures and tables.

## TECHNICAL VALIDATION

The sample used for this study contained 28 species (30 strains), including an archaeon (*Nitrososphaera viennensis* - Nv), one eukaryote (*Chlamydomonas reinhardtii* - CRH), 23 bacteria, and 5 bacteriophages (M13, P22, ES18, F0 and F2). Varying abundances of these species (at cell number and proteinaceous biomass levels) were used to create an uneven mock community (See **Supplementary Table 1** for details). Four biological replicates of the uneven mock community were processed using a sample preparation kit (see Methods) and submitted for ddaPASEF and diaPASEF analysis. The resulting ddaPASEF and diaPASEF data were processed using SpectroMine and Spectronaut, respectively. In both methods, the mass spectrometry data were searched against the protein sequence database (FASTA file) that contained sequences for all the organisms in the original ground-truth data set (see **Supplementary Table 1**) to generate peptide and protein outputs with quantitative values.

### Proteome depth

Database searches using the diaPASEF method detected greater numbers of precursors, peptides and proteins compared to those using the ddaPASEF method. In particular, 168% more precursors, 155% more peptides and 66% more proteins were detected using the diaPASEF method (**Figure 2A**). Although, it should be noted that 33% more peptide-digest was loaded for the diaPASEF method as compared to the ddaPASEF method (1 ug as compared to 750 ng). The overlap of detections amongst ddaPASEF and diaPASEF replicates demonstrates the reproducibility of the diaPASEF method with respect to the detection of precursors and proteins (**Supplementary Figure 1**).

**Figure 2:**
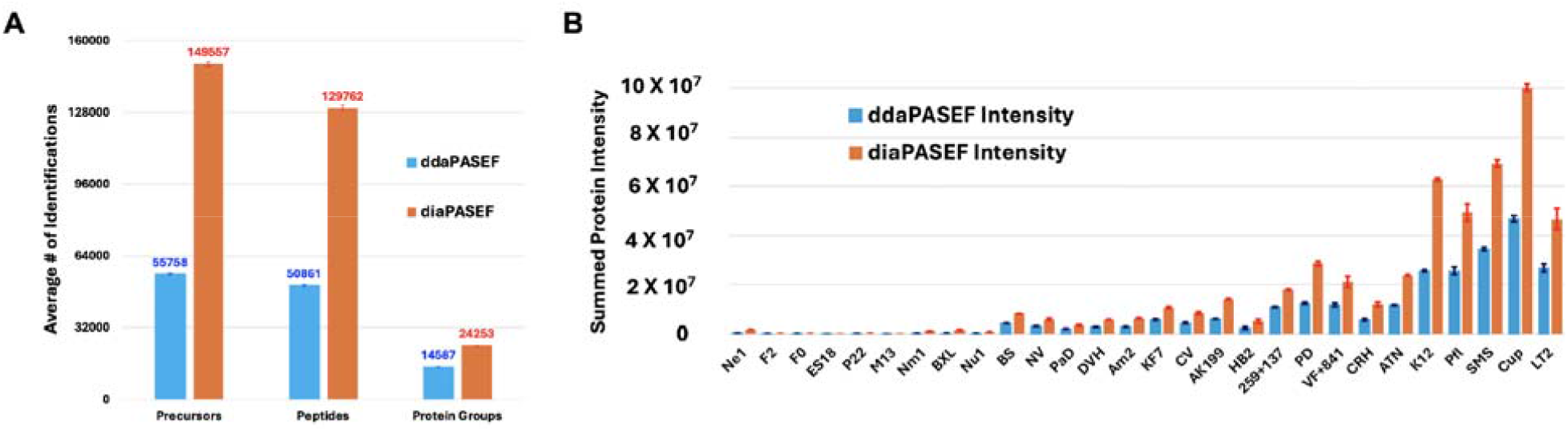
A) Comparison of identifications using ddaPASEF and diaPASEF mass spectrometry. The averages of identified peptide precursors, peptides, and protein groups were obtained from ddaPASEF (blue) and diaPASEF (orange) mass spectrometry methods. The peptide precursors, peptides and protein groups were detected at 1% FDR. SpectroMine and Spectronaut were used to search the ddaPASEF and diaPASEF data, respectively, against the protein sequence database that consisted of proteomes of the uneven mock community species. **B) Comparison of ddaPASEF and diaPASEF intensity measurements by organism**. Protein intensity measurements were obtained from ddaPASEF (blue) and diaPASEF (orange) searches using SpectroMine and Spectronaut, respectively. Organisms are ordered along the x-axis by protein abundance (smallest to largest).

When comparing protein intensities (**Figure 2B**) and protein groups (**Supplementary Figure 2A**) against the amount of protein input from each organism (**Supplementary Figure 2B** and **Supplementary Table 1)**, an expected positive correlation was observed between these measurements, wherein a greater protein input tended to correspond with greater protein intensity and number protein groups. However, there were exceptions to this observation: protein intensities (for Ne1, PD, CRH, K12 and LT2) and the number of protein groups (for Ne1, 199, HB2, VF and LT2) and did not demonstrate the expected trend of increasing abundance, as shown in **Figure 2B** and **Supplementary Figure 2A**, respectively.

### Quantitative reproducibility

To evaluate the reliability and consistency of quantitative measurements across acquisition modes, we assessed the coefficient of variation (CV) between replicates for ddaPASEF and diaPASEF datasets (**Figure 3**). This comparison aimed to determine which method provides more stable quantification across diverse microbial proteomes, particularly in complex metaproteomic samples. For 22 out of 28 organisms, diaPASEF yielded lower CV values than ddaPASEF - exceptions included F2, BS, KF7, 259, PD, and Cup. In the ddaPASEF dataset, four organisms (F0, M13, BXL, and Am2) exhibited CVs exceeding 20%, while in the diaPASEF dataset, only two (F2 and M13) surpassed this threshold. Notably, M13 showed extreme variability in both datasets, with CVs exceeding 170%, suggesting inconsistent detection of its proteomic features. Overall, the distribution of CV values (**Supplementary Figure 3**) indicates that diaPASEF delivers more reproducible quantification across biological replicates, reinforcing its suitability for comparative metaproteomics.

**Figure 3:**
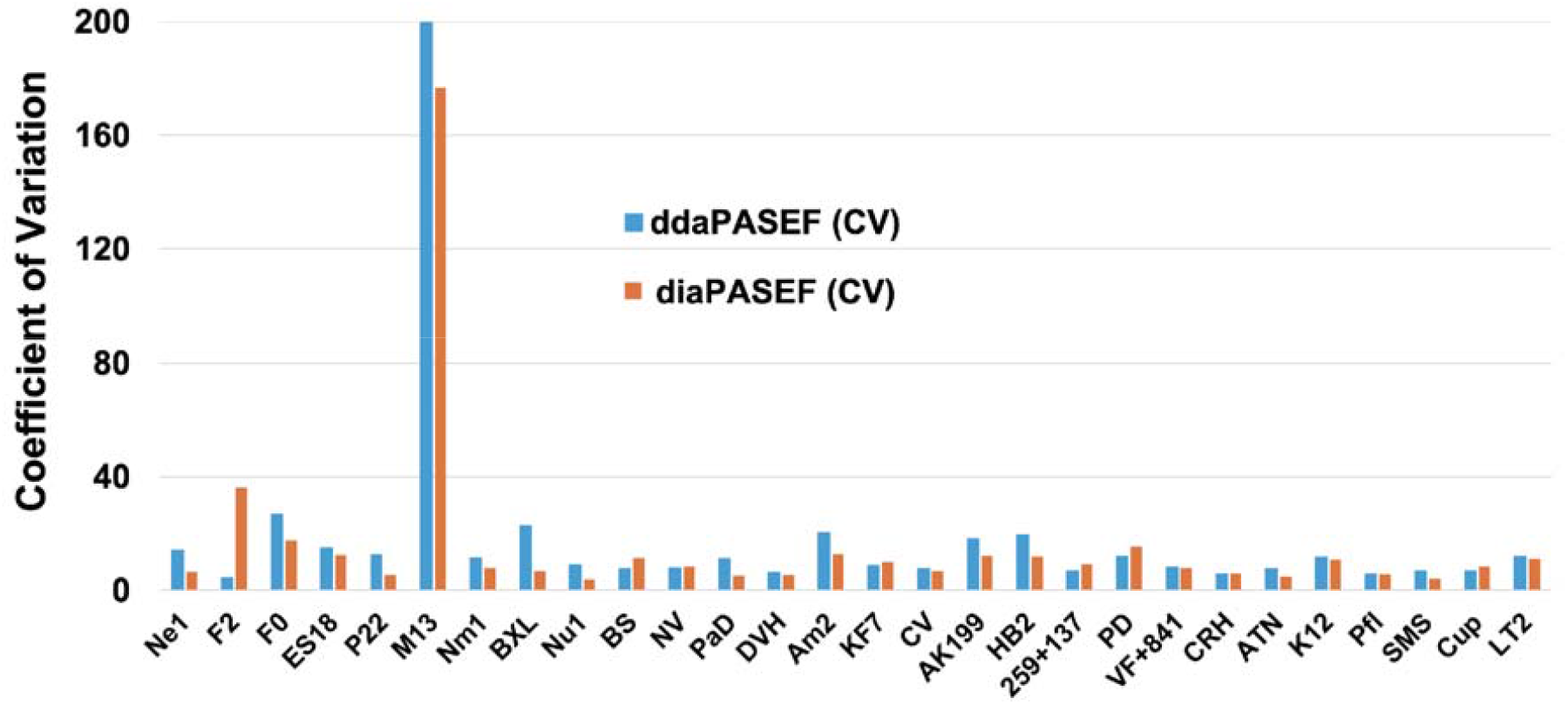
Comparison of coefficient of variation values for ddaPASEF and diaPASEF by organism. Coefficients of variation (CV) values were calculated based on intensity values for all proteins mapped to each organism and measured across replicates from ddaPASEF and diaPASEF searches. Most of the organisms had a coefficient of variation below 20% except F2 (diaPASEF data), F0 (ddaPASEF data), M13 (ddaPASEF and diaPASEF data) and BXL (ddaPASEF data) and Am2 (ddaPASEF data).

### Proteome coverage

Organism-level protein coverage averages were used to compare the performance of the two acquisition strategies. The resulting values were plotted as individual points for each organism, enabling direct visual comparison of coverage differences between ddaPASEF and diaPASEF (**Figure 4**). diaPASEF produced higher protein coverage for every species examined. This pattern highlights the increased depth and completeness of protein sequence information achievable with diaPASEF relative to ddaPASEF.

**Figure 4:**
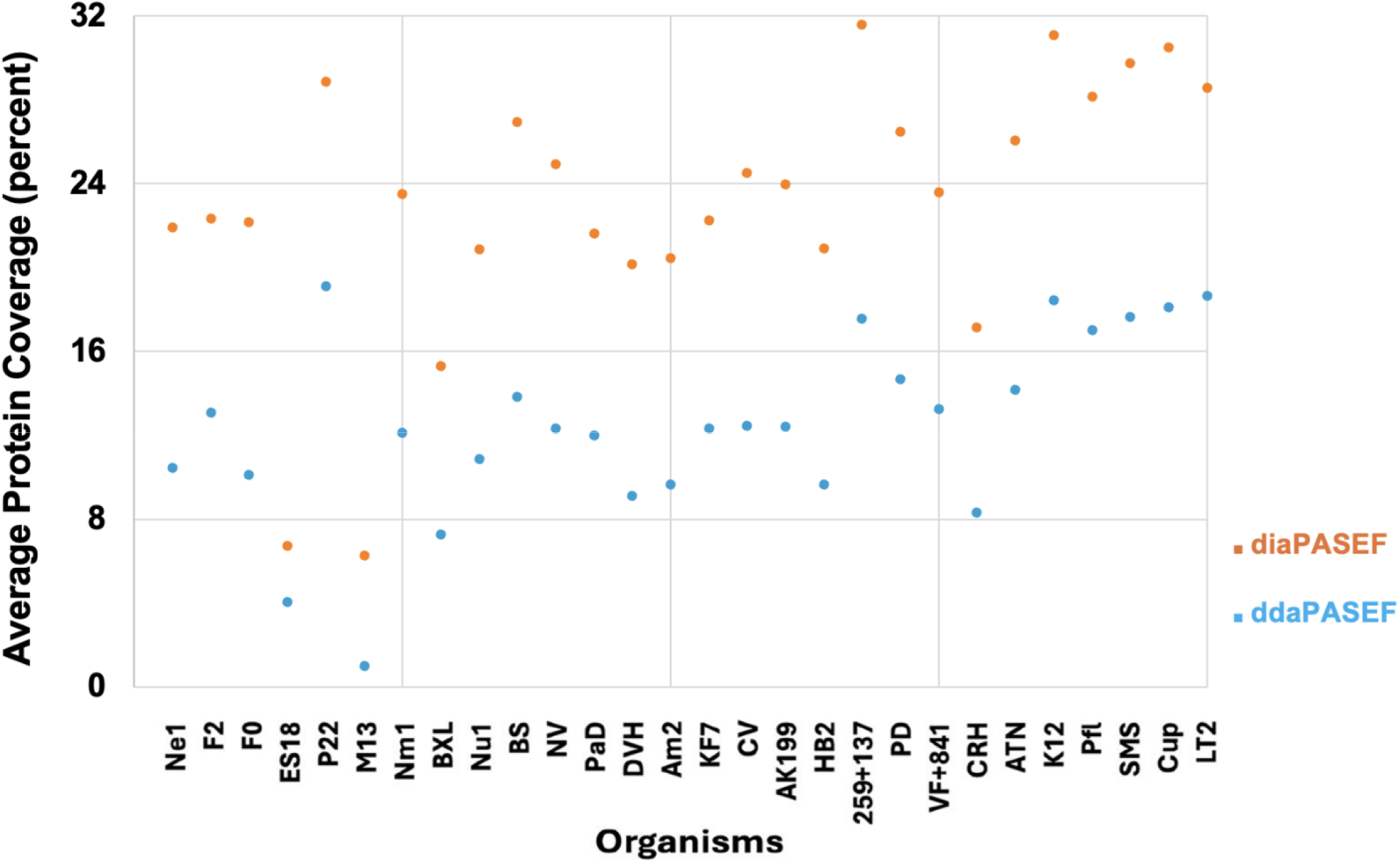
Protein coverage of species-specific proteins in ddaPASEF and diaPASEF. Average of protein coverage of species-specific proteins for each organism was calculated for both ddaPASEF data (blue) and diaPASEF data (orange). In all cases, the diaPASEF coverage was higher than ddaPASEF coverage.

### Quantitative accuracy

To examine the quantitative accuracy of the ddaPASEF and diaPASEF measurements, the measured percent abundances of each species were divided by their known percent abundances and the base-two log of the ratio calculated; the 95% confidence interval for these values is plotted in **Figure 5**, where the closer the interval is to zero the more accurate the measured values for that species are. We found that most of the organisms (the most abundant 22) showed that the log2(measured%/reference%) were close to zero for ddaPASEF and diaPASEF data (**Figure 5**). The reference refers to the expected relative abundance based on the composition of the ground truth community. However, the quantitative measurements of low-abundance bacteriophages were determined to have significant deviation from zero for the ddaPASEF and diaPASEF-dataset, with M13 showing both a high-variance and deviation from zero (**Figure 3**). The high variance for M13 can be explained due to a smaller number of M13 peptides and proteins detected (Table 1).

**Table 1:** Evidence for detection of five bacteriophages. The table shows the bacteriophage detection based on peptides and proteins per phage (in parentheses) in ddaPASEF and diaPASEF analysis of the uneven dataset.

|  | ddaPASEF |  |  |  | diaPASEF |  |  |  |
| --- | --- | --- | --- | --- | --- | --- | --- | --- |
| Phage | U1 | U2 | U3 | U4 | U1 | U2 | U3 | U4 |
| F2 | 1 (1) | 1 (1) | 1 (1) | 1 (1) | 1 (1) | 2 (1) | 2 (1) | 2 (1) |
| F0 | 4 (4) | 10 (6) | 7 (4) | 8 (4) | 26 (8) | 27 (8) | 27 (8) | 27 (8) |
| ES18 | 2 (2) | 3 (3) | 2 (2) | 2 (2) | 4 (3) | 4 (3) | 4 (3) | 4 (3) |
| P22 | 33 (7) | 38 (7) | 37 (7) | 32 (7) | 75 (8) | 71 (7) | 73 (8) | 75 (8) |
| M13 |  |  |  | 1 (1) |  | 1 (1) | 1 (1) | 3 (1) |

**Figure 5:**
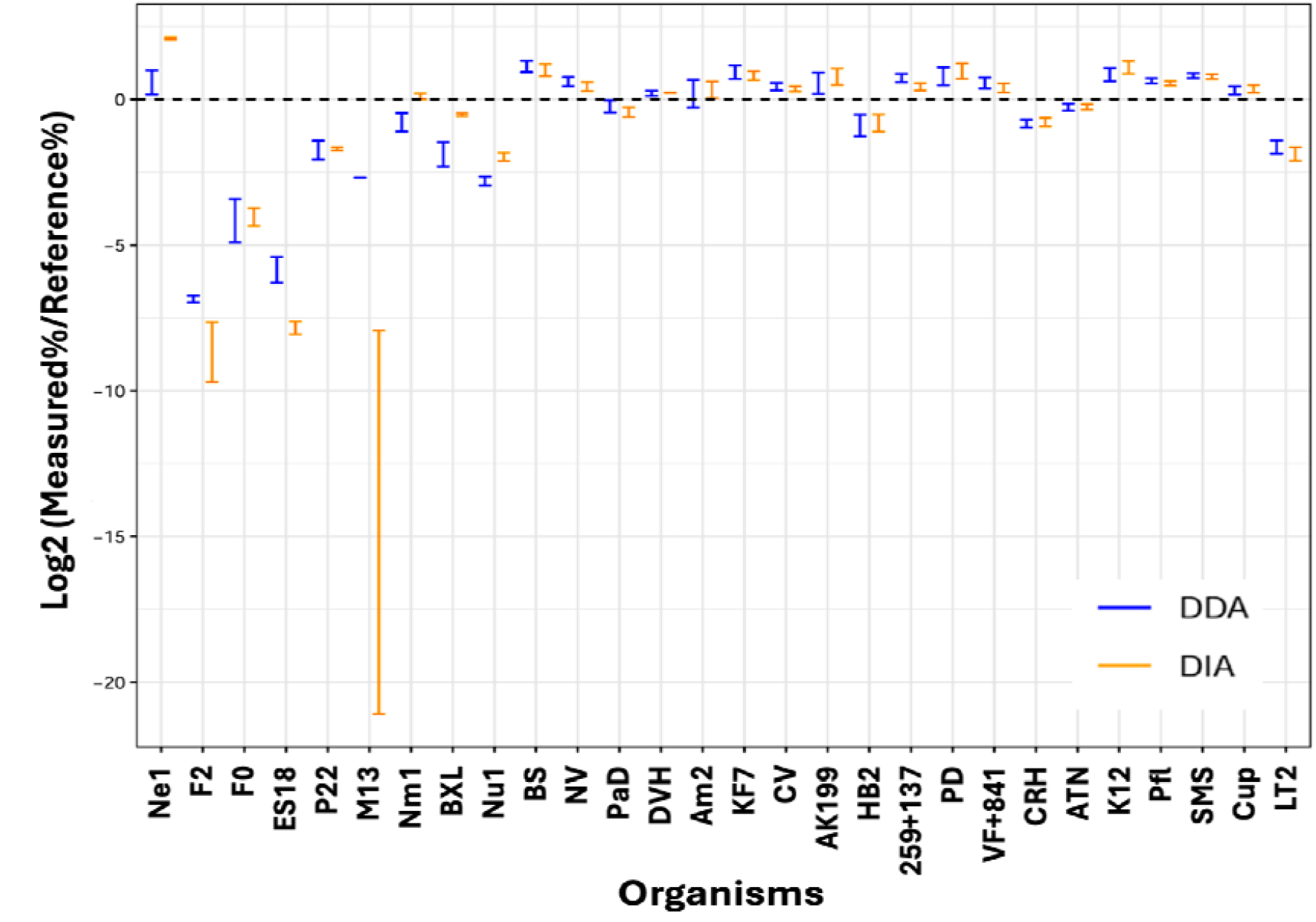
Quantitative accuracy of the mass spectrometry methods for determining species abundances in the uneven mock community. The 95% confidence interval of the log2(measured abundance/known abundance) was calculated to evaluate the quantitative accuracy of ddaPASEF and diaPASEF methods when determining species abundances of the uneven mock community. Organisms are ordered along the x-axis by abundance in the mock community (lowest to highest).

### Strain-level resolution

To evaluate the capability of both diaPASEF and ddaPASEF to differentiate between closely related microbial strains, we performed strain-associated variant (SAV) peptide analysis for *Staphylococcus aureus* (ATCC 13709 vs. ATCC 25923) and *Rhizobium leguminosarum* bv. *viciae* strains 3841 and VF39 (**Figure 6**).

**Figure 6:**
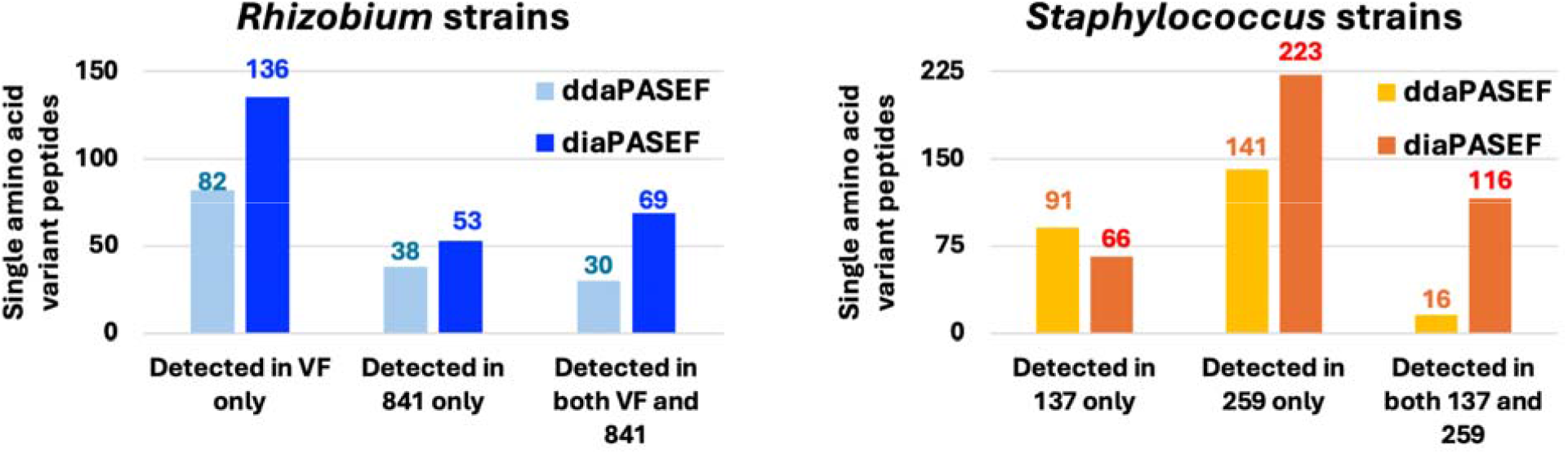
Detection of single amino acid variant peptides across bacterial strains using ddaPASEF and diaPASEF. Bar plots depict the detection of single–amino□acid□variant peptides detected across two bacterial genera - Rhizobium (left) and Staphylococcus (right) using two acquisition strategies, ddaPASEF and diaPASEF. For *Rhizobium*, peptides that were uniquely detected in strain VF or strain 841; or when SAV peptide-pair from both strains are detected, are shown. For *Staphylococcus*, peptides uniquely detected in strains 137 or 259, as well as peptides detected in both strains, are displayed. Across all categories and both strain pairs, diaPASEF consistently identifies a greater number of variant peptides than ddaPASEF, highlighting its improved sensitivity and coverage for strain□resolved proteomic variant detection.

For the *S. aureus* comparison, diaPASEF identified a greater number of strain-specific SAV peptides than the ddaPASEF dataset. A similar trend was observed for the *R. leguminosarum* bv. *viciae* strains 3841 and VF39. Interestingly, in cases where both the SAV peptide pairs were detected, diaPASEF detected more pairs (69 peptide-pairs in *Rhizobium* and 116 peptide pairs in *Staphylococcus*) than ddaPASEF (30 peptide-pairs in *Rhizobium* and 16 peptide pairs in *Staphylococcus*). Together, these findings highlight the discriminative power of the PASEF acquisition strategy for strain-level peptide detection and support the robustness of both diaPASEF and ddaPASEF workflows for reliable microbial strain identification.

### Bacteriophage detection

Since the protein intensities and quantitative accuracy for the bacteriophages were low, we decided to examine the detection evidence for the bacteriophages (**Table 1**). Bacteriophages were detected with more peptides (and sometimes proteins) using the diaPASEF analysis as compared to the ddaPASEF analysis. M13 was the only bacteriophage that was not detected in all replicates (in both ddaPASEF and diaPASEF analysis). We also investigated the distribution of Q-values for peptide detections via ddaPASEF or diaPASEF for three of the bacteriophages (M13, ES18 and F2) (**Supplementary Figure 4**). The Q-value is a statistical measure used to control the false discovery rate (FDR) in proteomics data analysis. It represents the minimum FDR at which a peptide appears in the filtered output list. A smaller Q-value provides higher confidence in the identification of peptides and proteins. In particular, when the Q-values from peptide identifications of the three bacteriophages across both ddaPASEF and diaPASEF datasets were ranked, the majority clustered within the lower Q-value range (**Supplementary Figure 4**), rather than being uniformly distributed or skewed toward higher Q-values. This pattern demonstrates that the bacteriophage-derived peptides were consistently detected at high confidence.

### Usage Notes

This dataset supports the a) Benchmarking of DIA and DDA acquisition strategies; b) Evaluation of protein inference algorithms; c) Development of strain-resolved metaproteomics workflows; d) Testing of mobility-aware scoring and feature detection; e) Assessment of quantitative reproducibility across complex communities; f) Validation of taxonomic assignment tools using ground-truth abundances and g) Training machine-learning models for peptide identification or quantification.

Our quantitative ground-truth dataset acquired using highly sensitive dda- and diaPASEF methods, provides a robust foundation for metaproteomics researchers to develop and rigorously evaluate metaproteomic bioinformatics methods. Beyond demonstrating the value of diaPASEF for deep and reproducible quantitative metaproteomics, this dataset will support the development of next□generation tools for more accurate peptide□to□protein mapping, improved assignment of peptides to microbial taxa and functions, and more precise quantification strategies tailored to complex community proteomes. It also creates an opportunity for emerging algorithms that leverage the ion□mobility dimension (including mobility□aware scoring, feature detection, and library construction) to be systematically tested and optimized. Together, these advances will help drive the creation of more specific, scalable, and biologically interpretable workflows for metaproteomics.

## Code Availability

Search parameter files, SpectroMine and Spectronaut configuration files are available in the MassIVE dataset folder.

## Data Availability

All raw and processed data are available at MassIVE under accession Reviewer access credentials are provided in the uploaded manuscript.

## Author Contributions

R.S. performed LC-MS acquisition and performed data analysis. A.T.R performed data analysis. A.O. and G.A. contributed to the computational analysis. K.D. and M.W. provided inputs during manuscript writing. M.K. provided mock community samples and biological validation. P.D.J. conceived and supervised the study, performed data analysis and wrote the manuscript. All authors contributed to manuscript preparation.

## CONFLICT OF INTEREST

R.S. and M.W. are employees of Bruker Daltonics GmbH & Co KG.

### ACKNOWLEDGEMENTS

Research reported in this publication was supported by the National Institute of General Medical Sciences of the National Institutes of Health (NIH) under Award Number R35GM138362 (MK). The content is solely the responsibility of the authors and does not necessarily represent the official views of the NIH. This research was supported in part by the Intramural Research Program of NIH. The contributions of the NIH authors are considered Works of the United States Government. The findings and conclusions presented in this paper are those of the authors and do not necessarily reflect the views of the NIH or the U.S. Department of Health and Human Services.

## DATA AVAILABILITY

Data can be accessed through MassIVE repository with an identifier **MSV000103357**. Dataset MSV000103357 is currently private.

For Reviewers: Log in into: https://massive.ucsd.edu/ProteoSAFe/static/massive.jsp using MSV000103357_reviewer as login and Metaproteomics1! as a password. Search for MSV000103357 dataset under “MassIVE Datasets” Tab. Once you have reached the dataset website for MSV000103357 you can use Browse Dataset Files to investigate the input files and search results.

For FTP access and dataset information use this link: ftp://MSV000103357@massive-ftp.ucsd.edu

We recommend using a dedicated FTP client to do this, though most web browsers will also support basic FTP access. Username for FTP access: **MSV000103357. Password for FTP access:** Metaproteomics1!

## REFERENCES

1. Sun Z, Ning Z, Figeys D. The Landscape and Perspectives of the Human Gut Metaproteomics. Mol Cell Proteomics. 2024 May;23(5):100763. doi: 10.1016/j.mcpro.2024.100763. Epub 2024 Apr 10. PMID: 38608842; PMCID: PMC11098955.

2. Zhao Z, Amano C, Reinthaler T, Baltar F, Orellana MV, Herndl GJ. Metaproteomic analysis decodes trophic interactions of microorganisms in the dark ocean. Nat Commun. 2024 Jul 30;15(1):6411. doi: 10.1038/s41467-024-50867-z. PMID: 39080340; PMCID: PMC11289388.

3. Van Den Bossche T, Lazau EA, Aho VTE, Alfredo Blakeley-Ruiz J, Wolf M, Kunath BJ, Valentin-Alvarado LE, Hellwig P, Verschaffelt P, Rajczewski AT, Henderickx JGE, Suomi T, Xian F, Shah S, Martens L, Benndorf D, Damare SR, Haak BW, Haange SB, Piehowski PD, Kupczok A, Figeys D, Mesuere B, Palmblad M, Hettich RL, Elo LL, Vizcaíno JA, Garg N, Wang Z, Arikan M, McCue LA, Griffin TJ, Clerbaux LA, Heyer R, Medema MH, Matallana-Surget S, Gomez-Varela D, Muth T, Tillett B, Armengaud J, Finn RD, Wilmes P, Gregory Caporaso J, Grenga L, Jagtap PD. Integrating multi-omics technologies to decipher microbiome functions. Nat Commun. 2026 Sep 8;17(1):9600. doi: 10.1038/s41467-026-77539-4. PMID: 42711342.

4. Van Den Bossche T, Armengaud J, Benndorf D, Blakeley-Ruiz JA, Brauer M, Cheng K, Creskey M, Figeys D, Grenga L, Griffin TJ, Henry C, Hettich RL, Holstein T, Jagtap PD, Jehmlich N, Kim J, Kleiner M, Kunath BJ, Malliet X, Martens L, Mehta S, Mesuere B, Ning Z, Tanca A, Uzzau S, Verschaffelt P, Wang J, Wilmes P, Zhang X, Zhang X, Li L; Metaproteomics Initiative. The microbiologist’s guide to metaproteomics. Imeta. 2025 May 6;4(3):e70031. doi: 10.1002/imt2.70031. PMID: 40469504; PMCID: PMC12130581.

5. Pietilä S, Suomi T, Elo LL. Introducing untargeted data-independent acquisition for metaproteomics of complex microbial samples. ISME Commun. 2022 Jun 29;2(1):51. doi: 10.1038/s43705-022-00137-0. PMID: 37938742; PMCID: PMC9723653.

6. Rajczewski AT, Blakeley-Ruiz JA, Meyer A, Vintila S, McIlvin MR, Van Den Bossche T, Searle BC, Griffin TJ, Saito MA, Kleiner M, Jagtap PD. Data-Independent Acquisition Mass Spectrometry as a Tool for Metaproteomics: Interlaboratory Comparison Using a Model Microbiome. Proteomics. 2025 May;25(9-10):e202400187. doi: 10.1002/pmic.202400187. Epub 2025 Apr 10. PMID: 40211604.

7. Gómez-Varela D, Xian F, Grundtner S, Sondermann JR, Carta G, Schmidt M. Increasing taxonomic and functional characterization of host-microbiome interactions by DIA-PASEF metaproteomics. Front Microbiol. 2023 Oct 16;14:1258703. doi: 10.3389/fmicb.2023.1258703. PMID: 37908546; PMCID: PMC10613666.

8. Wang A, Fekete EEF, Creskey M, Cheng K, Ning Z, Pfeifle A, Li X, Figeys D, Zhang X. Assessing fecal metaproteomics workflow and small protein recovery using DDA and DIA PASEF mass spectrometry. Microbiome Res Rep. 2024 Jul 3;3(3):39. doi: 10.20517/mrr.2024.21. PMID: 39421247; PMCID: PMC11480776.

9. Samodova D, Stankevic E, Søndergaard MS, Hu N, Ahluwalia TS, Witte DR, Belstrøm D, Lubberding AF, Jagtap PD, Hansen T, Deshmukh AS. Salivary proteomics and metaproteomics identifies distinct molecular and taxonomic signatures of type-2 diabetes. Microbiome. 2025 Jan 10;13(1):5. doi: 10.1186/s40168-024-01997-5. PMID: 39794871; PMCID: PMC11720885.

10. Xian F, Brenek M, Krisp C, Urbauer E, Ravi Kumar RK, Aguanno D, Srikumar T, Liu Q, Barry AM, Ma B, Krieger J, Haller D, Schmidt M, Gómez-Varela D. Ultra-sensitive metaproteomics redefines the dark metaproteome, uncovering host-microbiome interactions and drug targets in intestinal diseases. Nat Commun. 2025 Jul 18;16(1):6644. doi: 10.1038/s41467-025-61977-7. PMID: 40681571; PMCID: PMC12274446.

11. Xian F, Mitulovic G, Ravi Kumar RK, Uhrik L, Urbauer E, Aguanno D, Haller D, Schmidt M, Gomez-Varela D. Systematic evaluation of PASEF acquisition strategies in complex metaproteomes. Nat Commun. 2026 Aug 3;17(1):7406. doi: 10.1038/s41467-026-75845-5. PMID: 42547506; PMCID: PMC13434454.

12. Muth T, Kohrs F, Heyer R, Benndorf D, Rapp E, Reichl U, Martens L, Renard BY. MPA Portable: A Stand-Alone Software Package for Analyzing Metaproteome Samples on the Go. Anal Chem. 2018 Jan 2;90(1):685–689. doi:10.1021/acs.analchem.7b03544. Epub 2017 Dec 19. PMID: 29215871; PMCID: PMC5757220.

13. Li L, Ning Z, Cheng K, Zhang X, Simopoulos CMA, Figeys D. iMetaLab Suite: A one-stop toolset for metaproteomics. Imeta. 2022 May 21;1(2):e25. doi:10.1002/imt2.25. PMID: 38868572; PMCID: PMC10989937.

14. Alves G, Ogurtsov AY, Yu YK. Biological Function Assignment across Taxonomic Levels in Mass-Spectrometry-Based Metaproteomics via a Modified Expectation Maximization Algorithm. J Proteome Res. 2025 Aug 1;24(8):3818–3832. doi: 10.1021/acs.jproteome.4c01125. Epub 2025 Jul 18. PMID:40679470.

15. Vande Moortele T, Devlaminck B, Van de Vyver S, Van Den Bossche T, Martens L, Dawyndt P, Mesuere B, Verschaffelt P. Unipept in 2024: Expanding Metaproteomics Analysis with Support for Missed Cleavages and Semitryptic and Nontryptic Peptides. J Proteome Res. 2025 Feb 7;24(2):949–954. doi: 10.1021/acs.jproteome.4c00848. Epub 2025 Jan 10. PMID: 39792626.

16. Do K, Mehta S, Wagner R, Bhuming D, Rajczewski AT, Skubitz APN, Johnson JE, Griffin TJ, Jagtap PD. A novel clinical metaproteomics workflow enables bioinformatic analysis of host-microbe dynamics in disease. mSphere. 2024 Jun 25;9(6):e0079323. doi: 10.1128/msphere.00793-23. Epub 2024 May 23. PMID: 38780289; PMCID: PMC11332332.

17. Riffle M, May DH, Timmins-Schiffman E, Mikan MP, Jaschob D, Noble WS, Nunn BL. MetaGOmics: A Web-Based Tool for Peptide-Centric Functional and Taxonomic Analysis of Metaproteomics Data. Proteomes. 2017 Dec 27;6(1):2. doi: 10.3390/proteomes6010002. PMID: 29280960; PMCID: PMC5874761.

18. Sajulga R, Easterly C, Riffle M, Mesuere B, Muth T, Mehta S, Kumar P, Johnson J, Gruening BA, Schiebenhoefer H, Kolmeder CA, Fuchs S, Nunn BL, Rudney J, Griffin TJ, Jagtap PD. Survey of metaproteomics software tools for functional microbiome analysis. PLoS One. 2020 Nov 10;15(11):e0241503. doi: 10.1371/journal.pone.0241503. PMID: 33170893; PMCID: PMC7654790.

19. Schiebenhoefer H, Schallert K, Renard BY, Trappe K, Schmid E, Benndorf D, Riedel K, Muth T, Fuchs S. A complete and flexible workflow for metaproteomics data analysis based on MetaProteomeAnalyzer and Prophane. Nat Protoc. 2020 Oct;15(10):3212–3239. doi: 10.1038/s41596-020-0368-7. Epub 2020 Aug 28. PMID: 32859984.

20. Cheng, K., Figeys, D., MetaPilot enables adaptive genome-aware DDA and DIA metaproteomics using microbial genome catalogues. BioRxiv doi: 10.64898/2026.06.12.728088

21. Kleiner M, Thorson E, Sharp CE, Dong X, Liu D, Li C, Strous M. Assessing species biomass contributions in microbial communities via metaproteomics. Nat Commun. 2017 Nov 16;8(1):1558. doi: 10.1038/s41467-017-01544-x. PMID: 29146960; PMCID: PMC5691128.

22. Twigg CAI, Perez JM, Ryu J, Hanson BK, Barrera Estrada VJ, Thomas SN. Evaluation of Serum Proteome Sample Preparation Methods to Support Clinical Proteomics Applications. J Am Soc Mass Spectrom. 2024 Nov 6;35(11):2659–2669. doi: 10.1021/jasms.4c00131. Epub 2024 Sep 12. PMID: 39263706; PMCID: PMC11546599.

23. Zhang B, Chambers MC, Tabb DL. Proteomic parsimony through bipartite graph analysis improves accuracy and transparency. J Proteome Res. 2007 Sep;6(9):3549–57. doi: 10.1021/pr070230d. Epub 2007 Aug 4. PMID: 17676885; PMCID: PMC2810678.

