## Supplemental File for "Evaluation of Parallel Accumulation-Serial Fragmentation methods for metaproteomics using a model microbiome"

#### Supplementary Table 1

Protein sequence database components used in the proteomic analyses of composite microbiome samples.

| Label | Name and strain | Protein input per biological replicate (µg) | Protein abundance % |
|---|---|---|---|
| Ne1 | ***Nitrosomonas europaeae*** **ATCC 19718** | 60.58 | 0.082 |
| F2 | **Phage F2** | 62.04 | 0.084 |
| F0 | **Phage F0** | 65.28 | 0.088 |
| ES18 | **Phage ES18 (H1)** | 65.55 | 0.088 |
| P22 | **Phage P22 (HT105)** | 78.77 | 0.106 |
| M13 | **Phage M13** | 109.02 | 0.147 |
| Nm1 | ***Nitrosospira multiformis*** **ATCC 25196** | 155.16 | 0.209 |
| BXL | ***Burkholderia xenovorans*** **LB400** | 321.37 | 0.433 |
| Nu1 | ***Nitrosomonas ureae*** **Nm10** | 402.68 | 0.543 |
| BS | ***Bacillus subtilis*** **168** | 583.83 | 0.788 |
| Nv | ***Nitrososphaera viennensis*** | 607.41 | 0.819 |
| PaD | ***Paracoccus denitrificans*** **ATCC 17741, (Beijerinck and Minkman) Davis emends. Rainey et al.** | 683.66 | 0.922 |
| DVH | ***Desulfovibrio vulgaris*** **Hildenborough** | 701.58 | 0.946 |

| | | | |
|---|---|---|---|
| **Am2** | ***Alteromonas macleodii*** **ATCC 27126** | 707.47 | 0.954 |
| **KF7** | ***Pseudomonas pseudoalcaligenes*** **KF707** | 863.8 | 1.165 |
| **CV** | ***Chromobacterium violaceum*** **CV026** | 933.08 | 1.259 |
| **199** | ***Roseobacter*** **sp. AK199** | 1183.43 | 1.596 |
| **HB2** | ***Thermus thermophilus*** **HB27** | 1245.63 | 1.68 |
| **259 + 137** | ***Staphylococcus aureus*** **subsp.** ***aureus*** **Rosenbach, Seattle 1945, ATCC 25923;** ***Staphylococcus aureus subsp. aureus Rosenbach, Smith [NCTC 10399], ATCC 13709*** | 1931.46 | 2.606 |
| **PD** | ***Pseudomonas denitrificans*** **ATCC 13867** | 2128.51 | 2.871 |
| **VF + 841** | ***Rhizobium leguminosarum*** **bv.** ***viciae*** **VF39*****; Rhizobium leguminosarum bv. viciae 3841*** | 2351.5 | 3.172 |
| **CRH** | ***Chlamydomonas reinhardtii*** | 2962.73 | 3.996 |
| **ATN** | ***Agrobacterium tumefaciens*** **NTL4** | 4186.28 | 5.647 |
| **K12+ 2191** | ***Escherichia coli K12 with Flac+ Plasmid*** | 4290.54 | 5.788 |
| **Pfl** | ***Pseudomonas fluorescens*** **ATCC 13525, Type strain** | 4964.22 | 6.696 |
| **SMS** | ***Stenotrophomonas maltophilia*** **SeITE02** | 5946.27 | 8.021 |
| **Cup** | ***Cupriavidus metallireducens*** **CH34; (DSM 2839; LMG 1195; CIP 107179)** | 11504.98 | 15.519 |
| **LT2** | ***Salmonella enterica typhimurium*** **LT2** | 25037.78 | 33.774 |

**Supplementary Table 2**

The diaPASEF method covered the range of 300 to 1200m/z and an ion mobility range of 0.7-1.33 Vs cm−2, with 35Da isolation windows (1Da overlap) with a total cycle time of 850 ms. The collision energy (CE) was a linear function between the CE and the ion-mobility starting from 20 eV at 1/K0 = 0.6 Vs cm−2 to 59 eV at 1/K0 = 1.6 V cm−2. Accumulation and ramp time were set to 100 ms.

| #MS Type | Cycle Id | Start IM [1/K0] | End IM [1/K0] | Start Mass [m/z] | End Mass [m/z] | CE [eV] |
|---|---|---|---|---|---|---|
| **MS1** | 0 | - | - | - | - | - |
| **PASEF** | 1 | 0.7 | 0.865 | 404.5 | 440.5 | - |

| | | | | | | |
|---|---|---|---|---|---|---|
| **PASEF** | 1 | 0.865 | 1.055 | 649.5 | 685.5 | - |
| **PASEF** | 1 | 1.055 | 1.215 | 894.5 | 930.5 | - |
| **PASEF** | 1 | 1.215 | 1.3 | 1139.5 | 1200.5 | - |
| **PASEF** | 2 | 0.7 | 0.836 | 369.5 | 405.5 | - |
| **PASEF** | 2 | 0.836 | 1.029 | 614.5 | 650.5 | - |
| **PASEF** | 2 | 1.029 | 1.189 | 859.5 | 895.5 | - |
| **PASEF** | 2 | 1.189 | 1.3 | 1099.5 | 1140.5 | - |
| **PASEF** | 3 | 0.7 | 0.808 | 334.5 | 370.5 | - |
| **PASEF** | 3 | 0.808 | 1.005 | 579.5 | 615.5 | - |
| **PASEF** | 3 | 1.005 | 1.17 | 824.5 | 860.5 | - |
| **PASEF** | 3 | 1.17 | 1.3 | 1069.5 | 1100.5 | - |
| **PASEF** | 4 | 0.7 | 0.782 | 299.5 | 335.5 | - |
| **PASEF** | 4 | 0.782 | 0.964 | 544.5 | 580.5 | - |
| **PASEF** | 4 | 0.964 | 1.148 | 789.5 | 825.5 | - |
| **PASEF** | 4 | 1.148 | 1.3 | 1039.5 | 1070.5 | - |
| **PASEF** | 5 | 0.7 | 0.938 | 509.5 | 545.5 | - |
| **PASEF** | 5 | 0.938 | 1.126 | 754.5 | 790.5 | - |
| **PASEF** | 5 | 1.126 | 1.3 | 999.5 | 1040.5 | - |
| **PASEF** | 6 | 0.7 | 0.913 | 474.5 | 510.5 | - |
| **PASEF** | 6 | 0.913 | 1.101 | 719.5 | 755.5 | - |
| **PASEF** | 6 | 1.101 | 1.3 | 964.5 | 1000.5 | - |
| **PASEF** | 7 | 0.7 | 0.889 | 439.5 | 475.5 | - |
| **PASEF** | 7 | 0.889 | 1.078 | 684.5 | 720.5 | - |
| **PASEF** | 7 | 1.078 | 1.3 | 929.5 | 965.5 | - |

**Supplementary Table 3**

**Strain-associated variant (SAV) peptides detected by diaPASEF.** In cases where both the reference and the associated single amino acid variant (SAAV) were detected, peptide sequences identified experimentally are shown, with the amino acid variant shown in [ ]. Peptides originate from four bacterial strains analyzed in this study: *Staphylococcus aureus* ATCC 13709 (“137”), *S. aureus* ATCC 25923 (“259”), *Rhizobium leguminosarum* bv. *viciae* 3841 (“841”), and *R. leguminosarum* bv. *viciae* VF39 (“VF”). For peptides detected only in a single strain, please refer to the submission in the MassIVE repository.

| Peptide | Peptide SAV |
|---|---|
| **Identified in both *Rhizobium leguminosarum* bv. *viciae* 3481(“841”) and *Rhizobium leguminosarum* bv. *viciae VF39* (“VF”) strains** | |
| AEGSGDEA[A]MPIR | AEGSGDEA[E]MPIR |
| AGECHLIPYPNAADV[A]ELK | AGECHLIPYPNAADV[S]ELK |
| AGLFESV[D]VK | AGLFESV[E]VK |
| AISDLQAAGQNVSEQDV[A]AK | AISDLQAAGQNVSEQDV[S]AK |
| AL[N]AAIDAFLK | AL[D]AAIDAFLK |
| AQDATNANLS[A]ADIDALDK | AQDATNANLS[E]ADIDALDK |
| ASDVR[G]FTADQLK | ASDVR[A]FTADQLK |
| AVTPDA[D]HKYDNDFGNEWLK | AVTPDA[E]HKYDNDFGNEWLK |
| DAKGDPILIDSSFT[A]DPAVITR | DAKGDPILIDSSFT[P]DPAVITR |
| EAQPEAEQAQPEAKP[D]GGK | EAQPEAEQAQPEAKP[E]GGK |
| EGD[V]EISGPR | EGD[I]EISGPR |
| [L]HQNEASPVQK | [F]HQNEASPVQK |
| GDPILIDSSFT[A]DPAVITR | GDPILIDSSFT[P]DPAVITR |
| GGINE[A]AEAAADSQSMR | GGINE[V]AEAAADSQSMR |
| IFVTEF[A]DLMPK | IFVTEF[S]DLMPK |
| ILGYGL[D]EFR | ILGYGL[E]EFR |
| IT[D]ISETGGTFDK | IT[E]ISETGGTFDK |
| IVAEQENADYSG[V]IIR | IVAEQENADYSG[I]IIR |
| IVAEQENADYSG[V]IIRK | IVAEQENADYSG[I]IIRK |
| IVFEPNP[D]GDEFK | IVFEPNP[E]GDEFK |
| IVFEPNP[D]GDEFKFESK | IVFEPNP[E]GDEFKFESK |
| LLQAL[A]IESSASTFSTLK | LLQAL[S]IESSASTFSTLK |
| LSLPIWASSYEDPAV[A]K | LSLPIWASSYEDPAV[T]K |
| MLDPMVVGEEHY[D]VAR | MLDPMVVGEEHY[E]VAR |
| MLDPMVVGEEHY[D]VARK | MLDPMVVGEEHY[E]VARK |
| NIPIELAAPTQVAIG[G]GAASLNGLTLK | NIPIELAAPTQVAIG[D]GAASLNGLTLK |
| NLYDPSPNGYSTAVI[V]PR | NLYDPSPNGYSTAVI[M]PR |
| QIGELI[D]AYK | QIGELI[E]AYK |

| | |
|---|---|
| SDLAGI[V]ASNK | SDLAGI[M]ASNK |
| SEADLIE[G]WSTAR | SEADLIE[A]WSTAR |
| SIDAVYLPEMAFDL[D]AEAAR | SIDAVYLPEMAFDL[E]AEAAR |
| TGYAGSG[A]YK | TGYAGSG[P]YK |
| TVED[S]GLSGADYAPHGGSFPINVK | TVED[N]GLSGADYAPHGGSFPINVK |
| [A]SPEELER | [V]SPEELER |
| VTRV[D]VTDYQEF | VTRV[E]VTDYQEF |
| WPA[A]DVESLIR | WPA[E]DVESLIR |
| AEVA[G]VEAPTAEAR | AEVA[A]VEAPTAEAR |
| AIGYSDADLDALPQSV[Q]DGK | AIGYSDADLDALPQSV[E]DGK |
| ALASVPITLDVK[V]V | ALASVPITLDVK[I]V |
| AMNDVL[A]ALK | AMNDVL[T]ALK |
| ASMEYLTAR[V] | ASMEYLTAR[I] |
| AVVAPWDVL[V]FK | AVVAPWDVL[T]FK |
| [N]FNAAQTGAK | [D]FNAAQTGAK |
| DYV[A]AAKPAFEK | DYV[T]AAKPAFEK |
| EALHPYG[G]K | EALHPYG[S]K |
| EAQPAAP[D]AVTPTAESAKPEATPEAK | EAQPAAP[E]AVTPTAESAKPEATPEAK |
| EAQPAAP[D]AVTPTAESAKPEATPEAKPAAEAPAEK | EAQPAAP[E]AVTPTAESAKPEATPEAKPAAEAPAEK |
| ELAAQGYPT[A]GFR | ELAAQGYPT[V]GFR |
| GLNVGQSD[V]TSDYWQGDEGK | GLNVGQSD[I]TSDYWQGDEGK |
| GLNVGQSD[V]TSDYWQGDEGKFQK | GLNVGQSD[I]TSDYWQGDEGKFQK |
| IAATGTIDGQPIG[I]NGDIR | IAATGTIDGQPIG[M]NGDIR |
| IAVLLQDGESASDMSASAPAAAPAAAPQAAQEEKP[A]AATPASAPVPAEPK | IAVLLQDGESASDMSASAPAAAPAAAPQAAQEEKP[V]AATPASAPVPAEPK |
| IEPEVSAPPV[K] | IEPEVSAPPV[R] |
| I[V]EITPLTKPADIIAEIPR | I[L]EITPLTKPADIIAEIPR |
| LA[A]ALLDLAK | LA[S]ALLDLAK |
| LEVLT[G]NVK | LEVLT[S]NVK |
| LGLSITDDPLI[V]R | LGLSITDDPLI[L]R |
| LIEN[A]AAAGIPAVK | LIEN[T]AAAGIPAVK |
| LLTDGTEVDSGPATVPGT[P]TAEGK | LLTDGTEVDSGPATVPGT[L]TAEGK |
| REFIQ[D]NALSVANLDI | REFIQ[E]NALSVANLDI |
| [A]KQEPLTVFHSALENIAPHVEVR | [S]KQEPLTVFHSALENIAPHVEVR |
| SNHLLVDANLLQN[K] | SNHLLVDANLLQN[R] |
| [A]SFEDLGHDLGYALLLK | [S]SFEDLGHDLGYALLLK |
| SVAENIVYGAFDVV[Q]GK | SVAENIVYGAFDVV[H]GK |
| SVV[N]TVGGTPLK | SVV[D]TVGGTPLK |
| [A]YNWVFNKPEA | [T]YNWVFNKPEA |
| VAQETAPEADLVA[K] | VAQETAPEADLVA[R] |
| V[V]DPTGQPPGTIVVDTPSR | V[L]DPTGQPPGTIVVDTPSR |
| VLGVLEGIQ[K] | VLGVLEGIQ[R] |

| **Identified in both *Staphylococcus aureus* ATCC 13709 (“137”) and *Staphylococcus aureus* ATCC 25923 (“259”) strains** | |
|---|---|
| AEDN[P]FFPSPYSLSQYTAPK | AEDN[A]FFPSPYSLSQYTAPK |
| AFNEQPFA[I]VK | AFNEQPFA[V]VK |
| ALNSTIANV[D]EDIEK | ALNSTIANV[N]EDIEK |
| [V]NLGNQNIMAVAWYQNSAEAK | [A]NLGNQNIMAVAWYQNSAEAK |
| ATDAEN[L]EK | ATDAEN[V]EK |
| ATDAEN[L]EKEEAITK | ATDAEN[V]EKEEAITK |
| AVAGLSE[E]QLK | AVAGLSE[D]QLK |
| AYKAQGL[S]AWGF | AYKAQGL[G]AWGF |
| DGEVALFRP[E]ENFK | DGEVALFRP[D]ENFK |
| DIKDF[D]DVK | DIKDF[N]DVK |
| DVVVTFPEEYH[T]EELAGK | DVVVTFPEEYH[A]EELAGK |
| DVVVTFPEEYH[T]EELAGKEATFK | DVVVTFPEEYH[A]EELAGKEATFK |
| EEASANNLSDTSQEAQ[E]IQEAKK | EEASANNLSDTSQEAQ[A]IQEAKK |
| EFAQLNGEEIN[E]QPSNFK | EFAQLNGEEIN[D]QPSNFK |
| EIELEDPYENMG[T]K | EIELEDPYENMG[A]K |
| EIP[Q]LVTK | EIP[L]LVTK |
| EIVSEGTDFPDFDAAL[T]AYK | EIVSEGTDFPDFDAAL[A]AYK |
| EQV[D]QITAIFK | EQV[N]QITAIFK |
| EVDIDISNHTS[E]LIDNDILK | EVDIDISNHTS[D]LIDNDILK |
| FTELLV[E]K | FTELLV[D]K |
| GITTFDLVQ[D]K | GITTFDLVQ[N]K |
| GTFIIDPDGVVQASEINADGIGRDASTL[S]HK | GTFIIDPDGVVQASEINADGIGRDASTL[A]HK |
| GY[T]AEQMVEELSQDFTNISK | GY[S]AEQMVEELSQDFTNISK |
| IAITFSEE[E]NVDRIVQAK | IAITFSEE[G]NVDRIVQAK |
| IL[D]PNGYWNSTLR | IL[N]PNGYWNSTLR |
| IVIDFG[I]DK | IVIDFG[V]DK |
| KAVAGLSE[E]QLK | KAVAGLSE[D]QLK |
| LHLNDLHDSEISLISTSGTFSD[R] | LHLNDLHDSEISLISTSGTFSD[K] |
| LLE[S]AGLIK | LLE[A]AGLIK |
| LSVLG[T]DLLK | LSVLG[A]DLLK |
| MIAPVLEELA[T]DYEGK | MIAPVLEELA[A]DYEGK |
| NEGDVT[N]SNYNQNSAVK | NEGDVT[I]SNYNQNSAVK |
| NGEVSNIKPSLIN[I]K | NGEVSNIKPSLIN[V]K |
| [D]HLHLVFE | [N]HLHLVFE |
| NNKPQ[N]QPAAPK | NNKPQ[S]QPAAPK |
| NNMQEISSELQQSEQ[F]K | NNMQEISSELQQSEQ[S]K |
| NVE[Y]FTEMPVR | NVE[H]FTEMPVR |
| NVLSTLEQ[L]K | NVLSTLEQ[P]K |
| QSALQHNVEVN[D]QDELK | QSALQHNVEVN[N]QDELK |
| Q[N]FNDILNYGVQIK | Q[S]FNDILNYGVQIK |

| | |
|---|---|
| [T]ISYNQQNYDTIASGK | [S]ISYNQQNYDTIASGK |
| SNNGLSM[I]PWGTK | SNNGLSM[V]PWGTK |
| [T]VMYLLVNEGELSTFGPK | [S]VMYLLVNEGELSTFGPK |
| TAFDPNQSGN[I]FMAANFK | TAFDPNQSGN[T]FMAANFK |
| TEYHNLTF[T]PAQFETEIVQAK | TEYHNLTF[A]PAQFETEIVQAK |
| TGGLGASYSTSSNN[I]QVTTTMAPSSNGR | TGGLGASYSTSSNN[V]QVTTTMAPSSNGR |
| TNTYYVEATNN[N]PK | TNTYYVEATNN[S]PK |
| TTEVVGE[D]HVTGIR | TTEVVGE[N]HVTGIR |
| TVILPIV[E]R | TVILPIV[G]R |
| TYTFVFTDYVN[E]K | TYTFVFTDYVN[D]K |
| VEAH[L]INNMGHDHTR | VEAH[S]INNMGHDHTR |
| V[E]LPNGVELR | V[V]LPNGVELR |
| VYVPND[E]EGR | VYVPND[D]EGR |
| ADAQQN[K]FNK | ADAQQN[N]FNK |
| AE[S]QANQMVGDAVEK | AE[A]QANQMVGDAVEK |
| AFIDSQ[E]FK | AFIDSQ[G]FK |
| AFIETYQQQHP[E]DEVK | AFIETYQQQHP[D]DEVK |
| AHETAQN[T]DLK | AHETAQN[A]DLK |
| AQDSL[N]DNTTK | AQDSL[I]DNTTK |
| [V]QENGLTVVDAFNFEAPK | [A]QENGLTVVDAFNFEAPK |
| [T]SQDEVGSGVVYKK | [A]SQDEVGSGVVYKK |
| [S]VHLTEQGADK | [A]VHLTEQGADK |
| D[V]DFFNQAK | D[A]DFFNQAK |
| DAF[E]LPYTIIK | DAF[A]LPYTIIK |
| DA[T]IDMLGTASKVEVTK | DA[S]IDMLGTASKVEVTK |
| D[E]NAHPLTQGTFIK | D[G]NAHPLTQGTFIK |
| DITTIMDNGEAYGYAT[D]K | DITTIMDNGEAYGYAT[N]K |
| [E]NSEHFNVEIAK | [D]NSEHFNVEIAK |
| DQQSAFYEILNMPNLNE[E]QR | DQQSAFYEILNMPNLNE[A]QR |
| DVMT[Y]GGLK | DVMT[F]GGLK |
| EAGIYE[L]LLTVNK | EAGIYE[S]LLTVNK |
| EEASANNLSDTSQEAQEIQEAK[R] | EEASANNLSDTSQEAQEIQEAK[K] |
| EFIQQ[V]PSSPSYALFK | EFIQQ[A]PSSPSYALFK |
| EKGTEITS[I]FYPK | EKGTEITS[T]FYPK |
| ELG[T]RTFDEE | ELG[A]RTFDEE |
| FN[N]AIEASK | FN[S]AIEASK |
| FN[N]AIEASKVELTK | FN[S]AIEASKVELTK |
| FP[I]AVIEEVLK | FP[V]AVIEEVLK |
| GQFHEND[D]IEGVVVR | GQFHEND[V]IEGVVVR |
| GTEITS[I]FYPK | GTEITS[T]FYPK |
| HINSGELDGLE[N]EAAITK | HINSGELDGLE[S]EAAITK |

| | |
|---|---|
| IENQEGIDNI[E]EILEVSDGLMVAR | IENQEGIDNI[A]EILEVSDGLMVAR |
| IGDGD[I]EK | IGDGD[V]EK |
| IKVTT[L]DELTPLIGK | IKVTT[I]DELTPLIGK |
| IVNQQS[T]MPAITVALEK | IVNQQS[S]MPAITVALEK |
| KLDSSITIAD[Y]VVSPTLIHK | KLDSSITIAD[H]VVSPTLIHK |
| [E]LTQEQIEEAK | [K]LTQEQIEEAK |
| LGYDTGESETPITPVIIG[E]EK | LGYDTGESETPITPVIIG[D]EK |
| LNDSQAPKADAQQN[K]FNK | LNDSQAPKADAQQN[N]FNK |
| LN[E]VEQTNTPGSLNPK | LN[D]VEQTNTPGSLNPK |
| LTDELVN[V]FQK | LTDELVN[G]FQK |
| LTFSD[E]VVEK | LTFSD[D]VVEK |
| MAPQTAGEYEGT[H]YQFK | MAPQTAGEYEGT[N]YQFK |
| MFA[T]NVPQTTTHK | MFA[A]NVPQTTTHK |
| NKLTFSD[E]VVEK | NKLTFSD[D]VVEK |
| NQ[Q]ISYK | NQ[N]ISYK |
| PYDPN[H]PDNK | PYDPN[N]PDNK |
| QINPLDNEGLLAFGS[D]LQK | QINPLDNEGLLAFGS[G]LQK |
| SGEESEVLV[D]DK | SGEESEVLV[A]DK |
| SGEESEVLV[D]DKNK | SGEESEVLV[A]DKNK |
| SGTYDANINIADEMF[D]NK | SGTYDANINIADEMF[N]NK |
| SSA[E]VQQTQQASIPASQK | SSA[D]VQQTQQASIPASQK |
| TINGTLDLHDELE[E]TLAK | TINGTLDLHDELE[Q]TLAK |
| TP[T]DSGLEEYAEILSSK | TP[S]DSGLEEYAEILSSK |
| TTNDAN[N]IATNSELK | TTNDAN[S]IATNSELK |
| VAVVGQSGNLT[P]ADK | VAVVGQSGNLT[S]ADK |
| VDRGQQS[D]EDDLNAMK | VDRGQQS[G]EDDLNAMK |
| VDSTFSYS[L]GGK | VDSTFSYS[S]GGK |
| VDSTFSYS[L]GGKFDSTK | VDSTFSYS[S]GGKFDSTK |
| [I]FEQLSSK | [V]FEQLSSK |
| VVVIPTDEESMIARDVMT[Y]GGLK | VVVIPTDEESMIARDVMT[F]GGLK |
| YDQTFKGV[T]AK | YDQTFKGV[S]AK |
| YGINFNQ[T]LETGGVMLGK | YGINFNQ[A]LETGGVMLGK |
| YSD[Q]NKPYKEG | YSD[D]NKPYKEG |
| YVFSFKDPN[V]NGK | YVFSFKDPN[A]NGK |
| YVGD[V]AFLQIEPVEGELNYNK | YVGD[A]AFLQIEPVEGELNYNK |

#### Supplementary Table 4

**Strain-associated variant (SAV) peptides detected by ddaPASEF.** In cases where both the reference and the associated single amino acid variant (SAAV) were detected, peptide sequences identified experimentally are shown, with the amino acid variant shown in [ ]. Peptides originate from four bacterial strains analyzed in this study: *Staphylococcus aureus* ATCC 13709 ("137"), *S. aureus* ATCC 25923

(“259”), *Rhizobium leguminosarum* bv. *viciae* 3841 (“841”), and *R. leguminosarum* bv. *viciae* VF39 (“VF”). For peptides detected only in a single strain, please refer to the submission in the MassIVE repository.

| Peptide | Peptide SAV |
|---|---|
| **Identified in both *Rhizobium leguminosarum* bv. *viciae* 3481(“841”) and *Rhizobium leguminosarum* bv. *viciae VF39* (“VF”) strains** | |
| AEVA[G]VEAPTAEAR | AEVA[A]VEAPTAEAR |
| AMNDVL[A]ALK | AMNDVL[T]ALK |
| EAQPAAP[D]AVTPTAESAKPEATPEAKPAAEAPAEK | EAQPAAP[E]AVTPTAESAKPEATPEAKPAAEAPAEK |
| GGPFLAELDTFAGVAPDDPAVADL[K] | GGPFLAELDTFAGVAPDDPAVADL[R] |
| GLNVGQSD[V]TSDYWQGDEGK | GLNVGQSD[I]TSDYWQGDEGK |
| GLNVGQSD[V]TSDYWQGDEGKFQK | GLNVGQSD[I]TSDYWQGDEGKFQK |
| IAVLLQDGESASDMSASAPAAAPAAAPQAAQEEKP[A]AATPASAPVPAEPK | IAVLLQDGESASDMSASAPAAAPAAAPQAAQEEKP[V]AATPASAPVPAEPK |
| I[V]EITPLTKPADIIAEIPR | I[L]EITPLTKPADIIAEIPR |
| LAYISGQGGQD[G]TGALSPDFAVQVK | LAYISGQGGQD[S]TGALSPDFAVQVK |
| LLTDGTEVDSGPATVPGT[P]TAEGK | LLTDGTEVDSGPATVPGT[L]TAEGK |
| LMSV[D]DAIAEGAMALFGEK | LMSV[E]DAIAEGAMALFGEK |
| REFIQ[D]NALSVANLDI | REFIQ[E]NALSVANLDI |
| SVAENIVYGAFDVV[Q]GK | SVAENIVYGAFDVV[H]GK |
| VAQETAPEADLVA[K] | VAQETAPEADLVA[R] |
| VLGVLEGIQ[K] | VLGVLEGIQ[R] |
| AISDLQAAGQNVSEQDV[A]AK | AISDLQAAGQNVSEQDV[S]AK |
| AQDATNANLS[A]ADIDALDK | AQDATNANLS[E]ADIDALDK |
| A[S]TVGEINEAVMAAANGK | A[T]TVGEINEAVMAAANGK |
| DAKGDPILIDSSFT[A]DPAVITR | DAKGDPILIDSSFT[P]DPAVITR |
| GDPILIDSSFT[A]DPAVITR | GDPILIDSSFT[P]DPAVITR |
| [G]FTADQLKDELAK | [A]FTADQLKDELAK |
| GGINE[A]AEAAADSQSMR | GGINE[V]AEAAADSQSMR |
| IVFEPNP[D]GDEFKFESK | IVFEPNP[E]GDEFKFESK |
| MLDPMVVGEEHY[D]VAR | MLDPMVVGEEHY[E]VAR |
| MLDPMVVGEEHY[D]VARK | MLDPMVVGEEHY[E]VARK |
| NLYDPSPNGYSTAVI[V]PR | NLYDPSPNGYSTAVI[M]PR |
| RGAEGEEAAAEGEGAS[A]EAPVLADVASE | RGAEGEEAAAEGEGAS[T]EAPVLADVASE |
| SDLAGI[V]ASNK | SDLAGI[M]ASNK |
| SEADLIE[G]WSTAR | SEADLIE[A]WSTAR |
| TLDELNAYAH[A]K | TLDELNAYAH[T]K |
| **Identified in both *Staphylococcus aureus* ATCC 13709 (“137”) and *Staphylococcus aureus* ATCC 25923 (“259”) strains** | |
| EDSKE[E]QIK | EDSKE[A]QIK |
| EDSKE[E]QIKK | EDSKE[A]QIKK |
| E[I]QIKKSFAK | E[G]QIKKSFAK |

| | |
|---|---|
| EDSKE[I]QIK | EDSKE[A]QIK |
| EDSKE[I]QIKK | EDSKE[A]QIKK |
| EDSKE[E]QIK | EDSKE[G]QIK |
| EDSKE[I]QIK | EDSKE[G]QIK |
| EDSKE[E]QIKK | EDSKE[G]QIKK |
| EDSKE[I]QIKK | EDSKE[G]QIKK |
| EKQK[T]IYYK | EKQK[A]IYYK |
| IGD[F]FSK | IGD[L]FSK |
| MK[F]TAIAK | MK[L]TAIAK |
| MNKN[L]VIK | MNKN[I]VIK |
| QK[T]IYYK | QK[A]IYYK |
| SVLKS[E]R | SVLKS[D]R |
| SVLKS[E]R | SVLKS[N]R |

#### Supplementary Figure 1

**A**

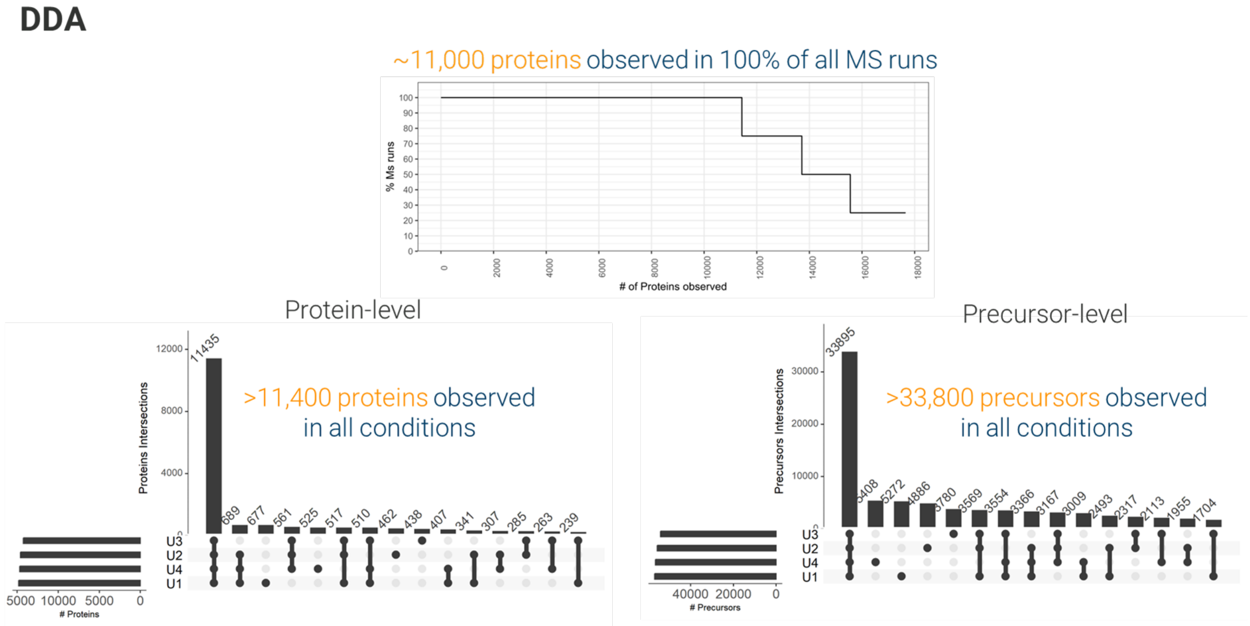


**B**

**DIA**

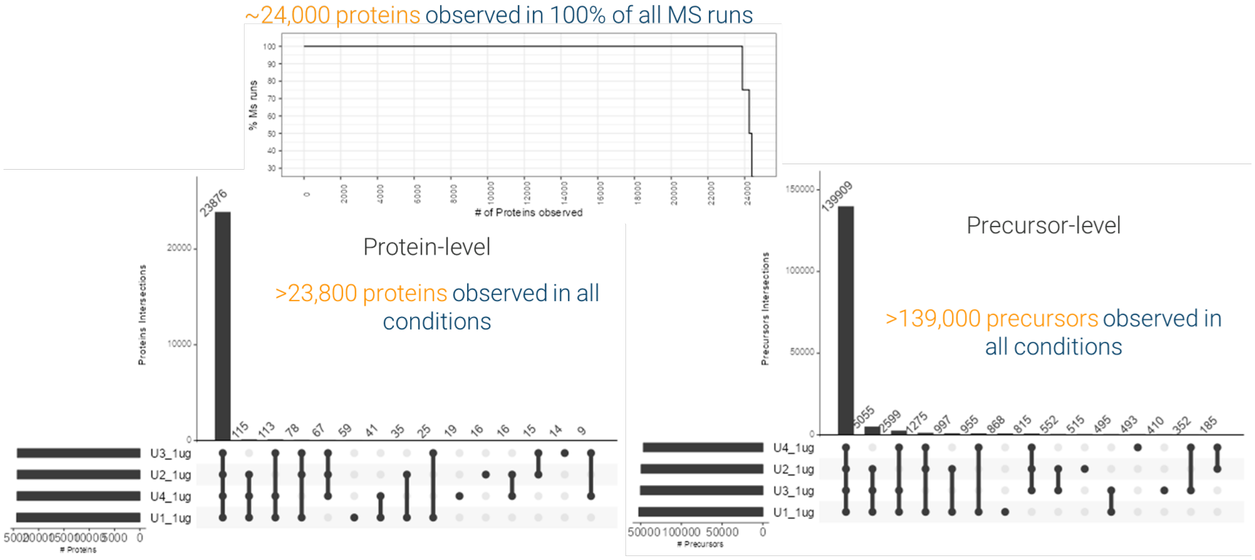
~24,000 proteins observed in 100% of all MS runs
% MS runs
### of Proteins observed
Protein-level
>23,800 proteins observed in all conditions
Proteins Intersections
23876
115
113
78
67
59
41
35
25
19
16
16
15
14
9
U3_1ug
U2_1ug
U4_1ug
U1_1ug
### Proteins
Precursor-level
>139,000 precursors observed in all conditions
Precursors Intersections
139909
5055
2599
1275
997
955
868
815
552
515
495
493
410
352
185
U4_1ug
U2_1ug
U3_1ug
U1_1ug
### Precursors

**C**

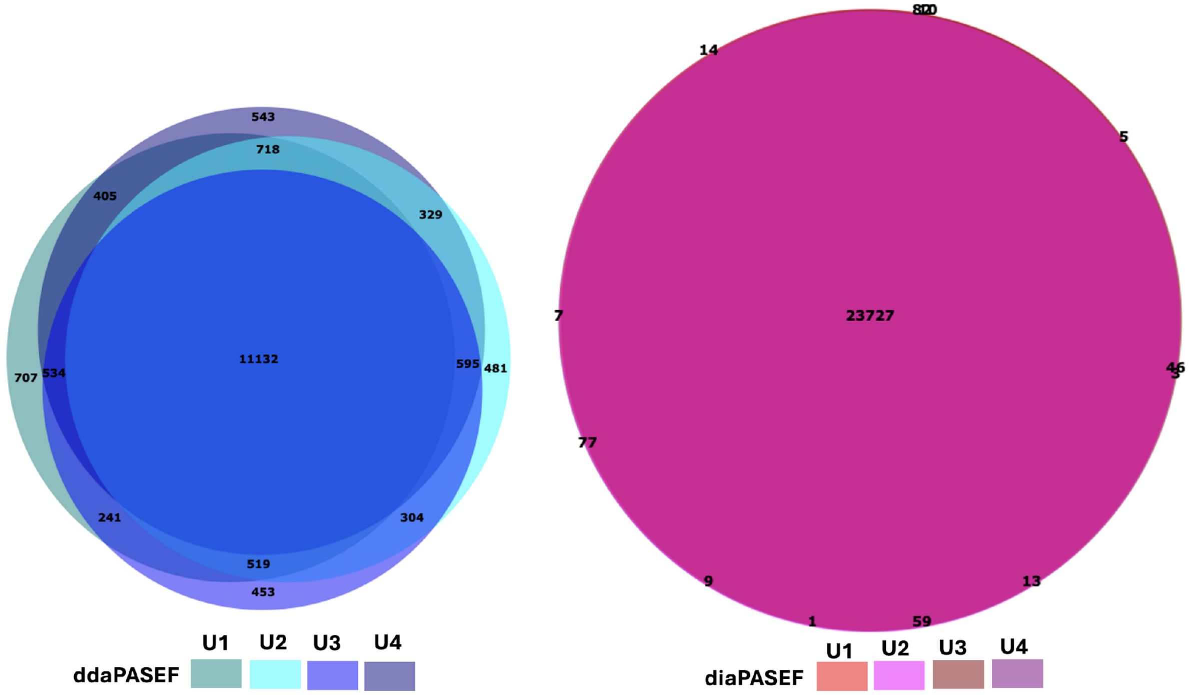
543
718
405
329
707
534
11132
595
481
241
304
519
453
U1
U2
U3
U4
ddaPASEF
14
5
7
23727
77
9
13
1
59
U1
U2
U3
U4
diaPASEF

**Supplementary Figure 1. Comparison of precursors and proteins** observed using A) ddaPASEF (DDA) and B) diaPASEF (DIA) methods across four replicates of uneven mock community samples (U1–U4). C) Venn diagram of proteins with quantitative values in ddaPASEF data and diaPASEF data.

**Supplementary Figure 2**

**A**

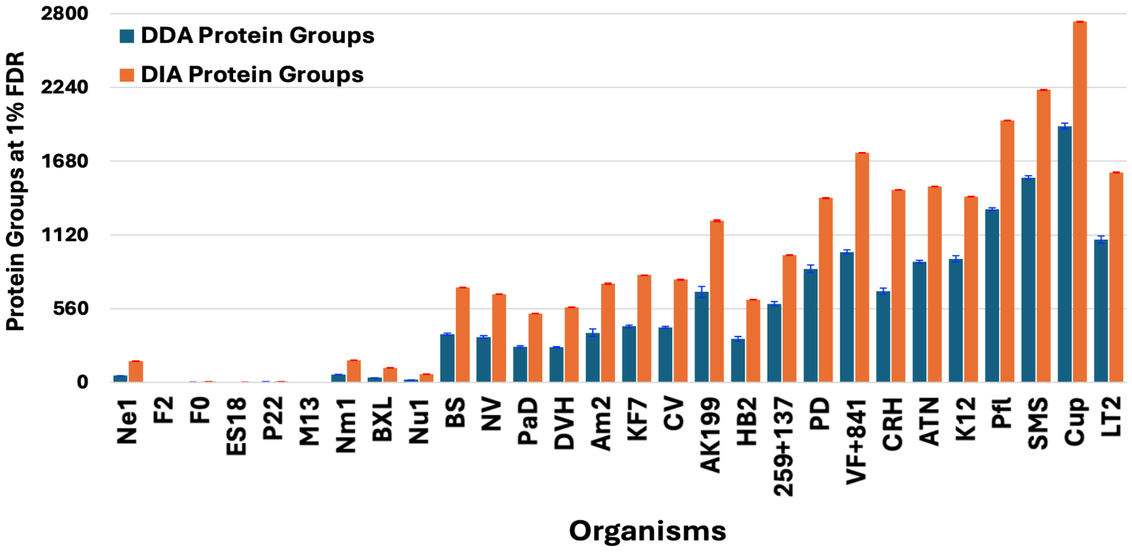


**B**

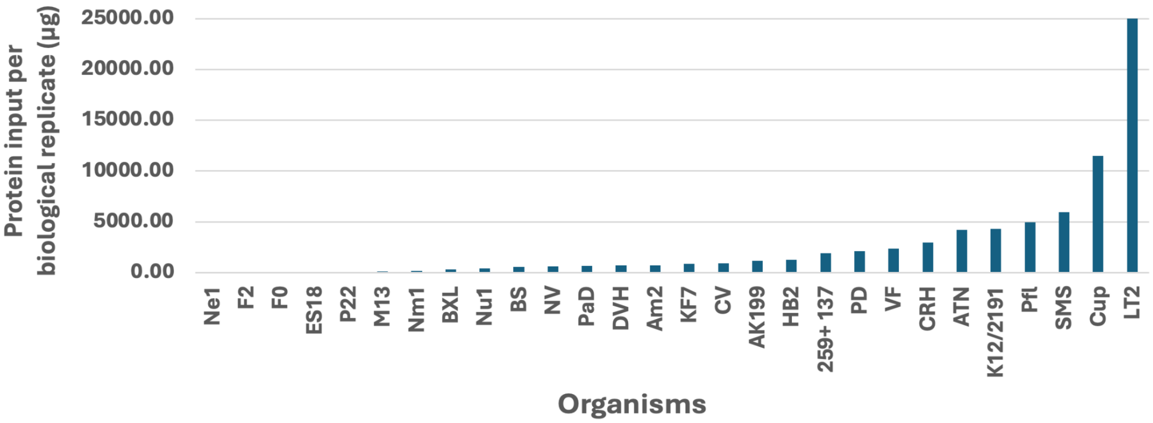


**Supplementary Figure 2: A) Comparison of ddaPASEF and diaPASEF protein groups by organism.** Average protein groups were obtained from ddaPASEF (DDA, blue) and diaPASEF (DIA, orange) searches using SpectroMine and Spectronaut, respectively. Organisms were ordered along the x-axis by protein abundance (smallest to largest). **B) Comparison of protein input per biological replicate by organism.** Organisms are ordered along the x-axis by protein abundance (smallest to largest). See **Supplementary Table 1** for organism names and strains.

**Supplementary Figure 3**

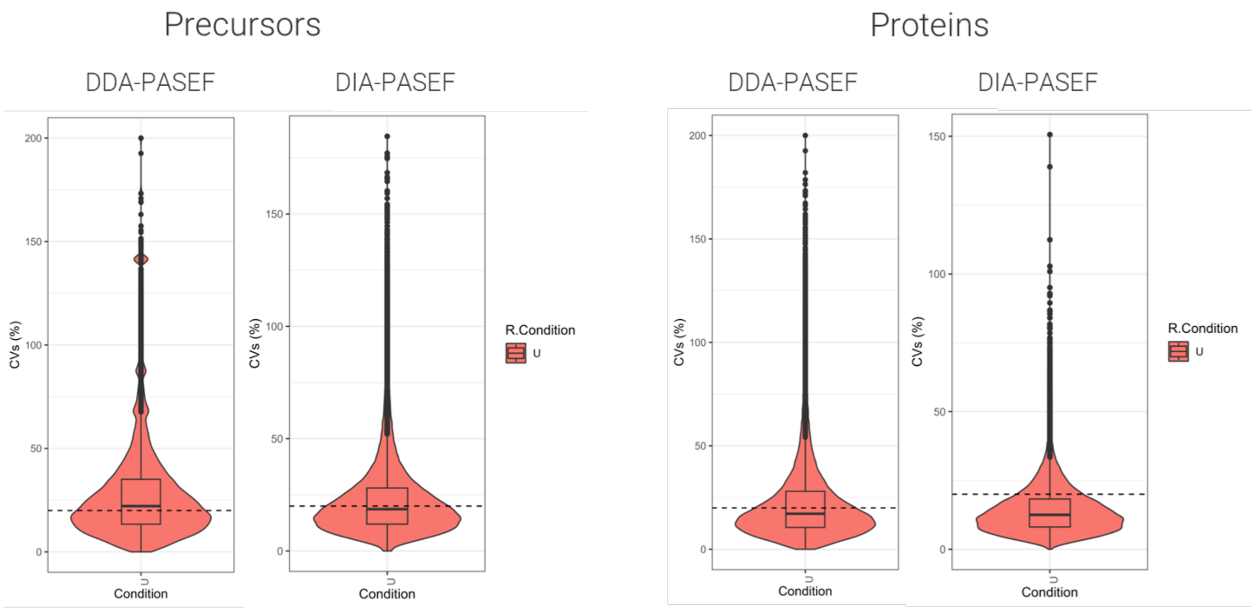


**Supplementary Figure 3. Distribution of ddaPASEF and diaPASEF CV values.** The distribution of CVs for ddaPASEF (DDA) and diaPASEF (DIA) methods were calculated at the precursor-level (left) and protein-level (right). The distribution of CV values for diaPASEF data sets indicates that the diaPASEF dataset has lower variances than those of the ddaPASEF dataset.

**Supplementary Figure 4**

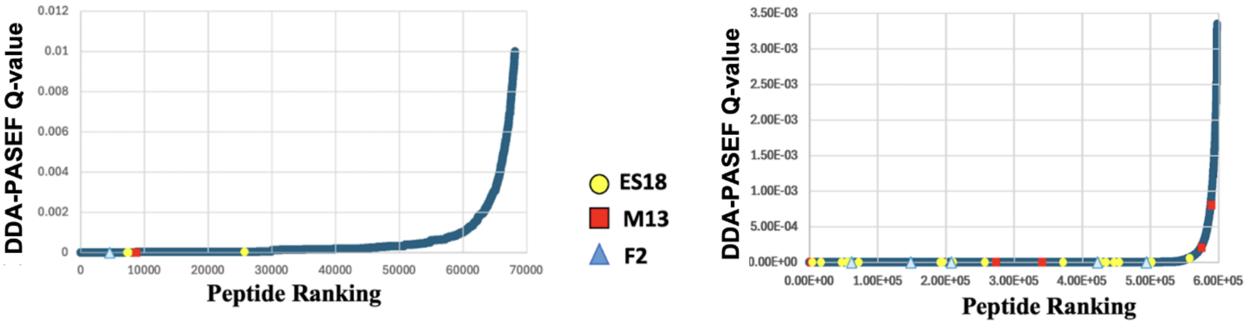


**Supplementary Figure 4: Evidence for detection of five bacteriophages using Q-values.** Q-values from peptide results for Spectromine (ddaPASEF) and Spectronaut (diaPASEF) were used for bacteriophages F2, ES18, and M13 to assess the confidence of their detection.
